# TNFPred: Identifying tumor necrosis factors using hybrid features based on word embeddings

**DOI:** 10.1101/860791

**Authors:** Trinh-Trung-Duong Nguyen, Nguyen-Quoc-Khanh Le, Quang-Thai Ho, Dinh-Van Phan, Yu-Yen Ou

## Abstract

**Background:** Cytokines are a class of small proteins that act as chemical messengers and play a significant role in essential cellular processes including immunity regulation, hematopoiesis, and inflammation. As one important family of cytokines, tumor necrosis factors have association with the regulation of a various biological processes such as proliferation and differentiation of cells, apoptosis, lipid metabolism, and coagulation. The implication of these cytokines can also be seen in various diseases such as insulin resistance, autoimmune diseases, and cancer. Considering the interdependence between this kind of cytokine and others, classifying tumor necrosis factors from other cytokines is a challenge for biological scientists. In this research, we employed a word embedding technique to create hybrid features which was proved to efficiently identify tumor necrosis factors given cytokine sequences. We segmented each protein sequence into protein words and created corresponding word embedding for each word. Then, word embedding-based vector for each sequence was created and input into machine learning classification models. When extracting feature sets, we not only diversified segmentation sizes of protein sequence but also conducted different combinations among split grams to find the best features which generated the optimal prediction. Furthermore, our methodology follows Chou’s 5-step rules to build a reliable classification tool.

**Results:** With our proposed hybrid features, prediction models obtain more promising performance compared to seven prominent sequenced-based feature kinds. Results from 10 independent runs on the surveyed dataset show that on an average, our optimal models obtain an area under the curve of 0.984 and 0.998 on 5-fold cross-validation and independent test, respectively.

**Conclusions:** These results show that biologists can use our model to identify tumor necrosis factors from other cytokines efficiently. Moreover, this study proves that natural language processing techniques can be applied reasonably to help biologists solve bioinformatics problems efficiently.

## Introduction

Cytokines is a varied group of polypeptides, usually linked to inflammation and cell differentiation or death. Among major families of cytokines (interleukins (IL), interferons (IFNs), tumor necrosis factors (TNFs), chemokine and various growth factors, comprised of transforminggrowth factor b(TGF-b), fibroblast growth factor (FGF), heparin binding growth factor (HBGF) and neuron growth factor (NGF)) [1], tumor necrosis factors are versatile cytokines with a wide range of functions that attracts abundant of biological researchers (see, e.g. [2-6]). TNFs can take part in pathological reactions as well as involve in a variety of processes, such as inflammation, tumor growth, transplant rejection, etc. [3, 6]. TNFs act through their receptors at the cellular level to activate separate signals that control cell survival, proliferation or death. Furthermore, TNFs play two opposite roles in regard to cancer. On the positive side, activity in the suppression of cancer is supposed to be limited, primarily due to system toxicity of TNFs. On the negative side, TNFs might act as a promoter of the endogenous tumor through their intervention to the proliferation, invasion and tumor cell metastasis thus contributing to tumor provenance. Such TNFs’ effect on cancer cell death makes them a probable therapeutic for cancer [3]. Moreover, in the United States and other nations, patients with TNF-linked autoimmune diseases have been authorized to be treated with TNF blockers [2]. In cytokine network, TNFs and other factors such as interleukins, interferons form an extremely complicated interactions generally mirroring cytokine cascades which begin with one cytokine causing one or additional different cytokines to express that successively trigger the expression of other factors and generate complex feedback regulatory circuits. Abnormalities in these cytokines, their receptors, and the signaling pathways that they initiate involve a broad range of illnesses [7-12]. Interdependence between TNFs and other cytokines accounts for such diseases. For instance, TNFs and interleukin-1 administers TNF-dependent control of mycobacterium tuberculosis infection [12]. Another example is the TNF and type I interferons interactions in inflammation process which involve rheumatoid arthritis and systemic lupus erythematosus [13]. For the above reasons, identification of TNFs from other cytokines presents a challenge for many biologists.

To date, in bioinformatics, several research teams have built machine learning models to predict cytokines and achieved high performance [14-20]. Bases on these research, it is noted that Support Vector Machine (SVM) [21] classifier is a solid foundation for building prediction models. Regarding the feature extraction method used, considerable efforts were made to create hybrid features through useful characteristics extracted from sequences such as compositions of amino acids and amino acid pairs, physicochemical properties, secondary structures, and evolutionary information to improve predictive ability. It therefore has been verified that a primary concern for biological scientists is a discriminatory and effective feature set. Additionally, among these groups, some have further classified cytokines into tumor necrosis factor family and other subfamilies. For example, Huang et al. utilized dipeptide composition to developed CTKPred, a tool for identifying cytokines families and subfamilies [15]. This classification method accomplished an accuracy of 92.5% on 7-fold cross-validation when predicting cytokines, and also enabled 7 significant cytokine classes to be predicted with 94.7% accuracy, overall. Although the results seem promising, we noted that the authors set cutoff threshold value as very high as 90% for excluding homologous sequences within the dataset. Three years later, Lata et al. constructed CytoPred using the combination of support vector machine and Psi Blast to classify cytokines into 4 families and 7 subfamilies [16]. Despite the fact that CytoPred outperforms CTKPred, Lata’s group reused the dataset created by Huang et al. which poses the similar concern about the high value for cutoff threshold. We believe that if the cutoff threshold is lower, the more the homology bias will be excluded from the surveyed dataset in a stricter manner and thus increasing reliability of the prediction model [22]. Considering the important roles of TNFs and the imperfection of such models like CTKPred and CytoPred, a fresh approach to classifying TNF among cytokines is required, so our study seeks to discover a solution to this issue.

From the interpretation of the genetic alphabet order, scientists have found that in terms of in composition, biological sequences, particularly protein sequences, are comparable to human language [23]. A growing variety of scientists are relating these molecule sequences as a special textual data and examining them using available text mining techniques. Initial stage of this transformation is to convert (biological) words to real numbers like those in vectors. This means that every word is encoded by one or more values that locate it in a discretionary space. Regarding this issue, NLP researchers have seen some landmarks. They are one-hot encoding, co-occurrence matrix, and word embedding techniques. A one-hot vector represents a word in text and encompass a dimension up to a vocabulary size, where a sole entry resembling the word is a one and all opposite entries are zeros. This presents a significant drawback because this method does not group commonly co-occurring items together in the representation space. Furthermore, scalability is another limitation as the representation size increases with the corpus size. Later, the concept of co-occurrence matrix arrived. Co-occurrence matrix represents a word while considering its adjacent words. The underlying reasoning for co-occurrence matrix follows an assumption that “context is king”. This technique has been believed to take a radical change within the field of NLP. During this methodology, a window size s is selected to formulate a co-occurrence matrix where words appearing along in the chunk with length s are going to be counted. The linguistics and grammar relationships made by this method are compelling yet this idea encounters some limitations such as high-dimensional space and thus computational expensive. These barriers resulted in the arrival of word embeddings, or continuous word vectors that are illustrations of words that also considers the context via neighboring words yet keep the number of dimensions greatly lower. A typical word embedding vector accumulates additional characteristics with reduced dimensions, and therefore more efficient. Recently, its utilization have been seen as underlying principle of various word embedding-related research such as sentiment classification, bilingual word translation, information retrieval, etc. with appreciable accomplishment [24-27].

Motivated by remarkable achievements in NLP using word embeddings, in this study, we tried to use NLP technique for extracting features. We transferred the protein sequences into “protein sentences” comprised of composing biological words from which word vectors were created. Next, we trained the Fast Text model to created word embeddings on which final word embedding-based features were generated. Finally, we employed some advanced machine learning algorithms for classification.

Many studies in bioinformatics show that [28-37] to build a helpful statistical predictor from primary sequences to help biologists solve a certain problem, researchers ought to obey the rules from 5-step rule which is restated here for clarity: (1) the accurate procedure to compose training and independent dataset to build and evaluate the predictive model; (2) the way to represent amino acid chain samples in mathematical forms that can genuinely mirror their characteristic interconnection with the predicted target; (3) the best approach to present or build up an effective and robust predictive algorithm; (4) the method to correctly conduct cross-validation tests to justly evaluate the predictor’s unseen accuracy; (5) the way to set up a convenient and accessible webserver for the predictor that is helpful to general users. Herein, we would strictly follow all the above-mentioned steps.

## Methodology

We present a new method utilizing word embedding vectors to efficiently identify tumor necrosis factors from cytokines. Figure1 displays a flowchart of our study consisting two main subprocesses: using FastText to train vector model and support vector machine classifier to train supervised learning classification. Each sub-process is explained further in the sections below.

**Figure 1.**
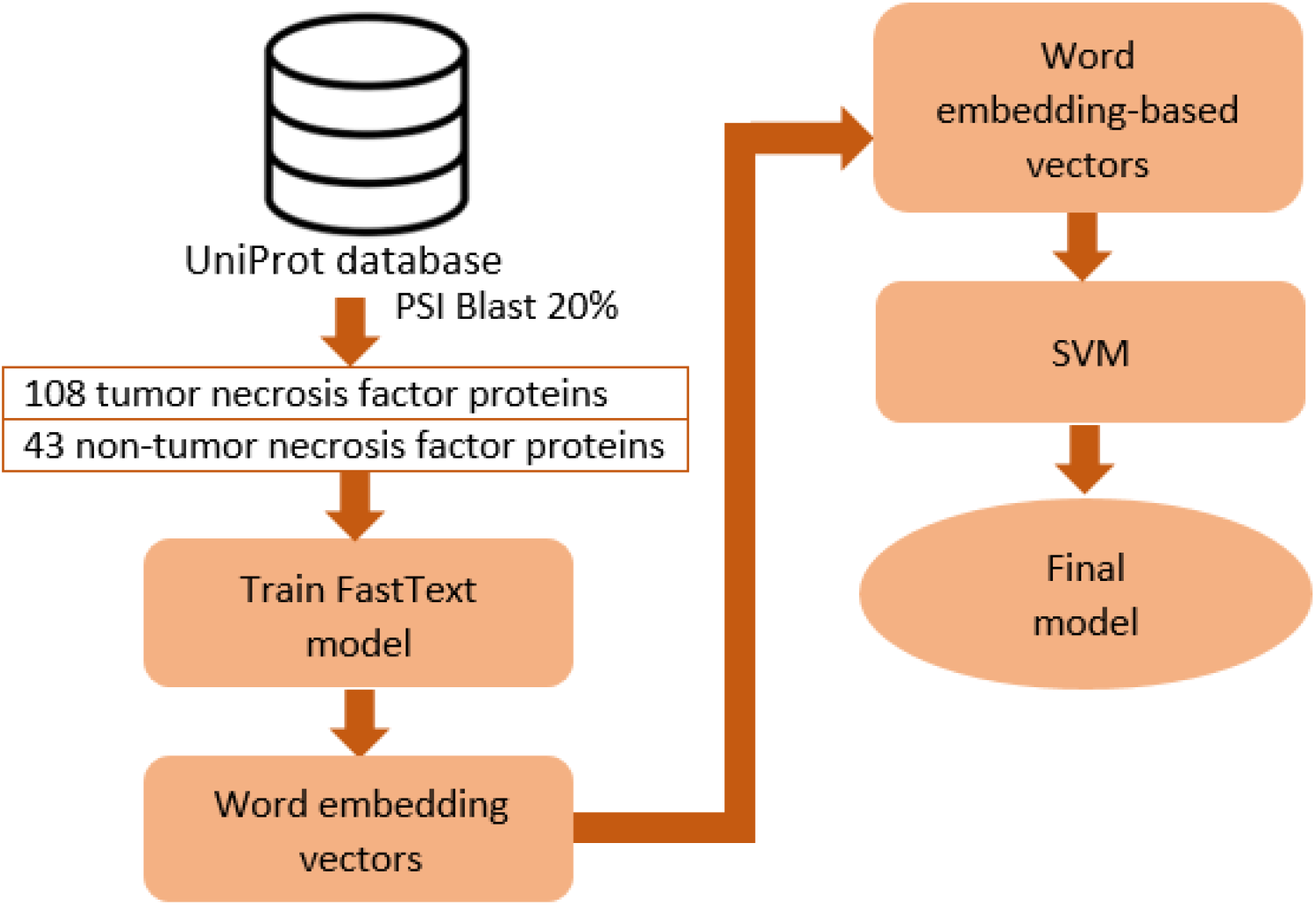
The flowchart of this study. First, surveyed dataset was used to train a FastText model and then we used this trained model to generate word embedding vectors. Next, word embedding-based vectors were created. In the end, support vector machine classifier was used for classification

### Data collection

From UniProt database [38] (release 2019_05), we first collected positive data which are tumor necrosis factors by using the query as “family:”tumor necrosis factor family” AND reviewed:yes”. From this step, we obtained 106 protein sequences. After that, we collected negative data which are 1023 cytokine sequences from other major cytokine families including 347 interleukins, 205 chemokines, 227 TGF-betas, 138 interferons and 106 others. Using PSI Blast [39], we dismissed sequences with similarity higher than 20% which ensures our study strictly follows the first principle in the 5-step rule. We also removed protein sequences that contain uncommon amino acid (BJOUXZ). After this step, we were left with 18 TNF proteins and 133 non-TNF proteins. These are proteins from various organisms such as human, mouse, rat, bovine and fruit fly, etc. As the number of protein sequences left for survey is small (151 sequences), the prediction model might be biased toward the way we separated the data parts used for training and testing. Accordingly, we randomly divided 151 surveyed sequences into cross-validation data and independent data for building and testing the models, respectively. We repeated this with process for 10 times when keeping the same sequence number distributions over these two parts. This means that all our later experiments were carried out on 10 different datasets and the results were averaged. Table 1 shows the detailed number of sequences for each part in each of 10 datasets.

**Table 1.**
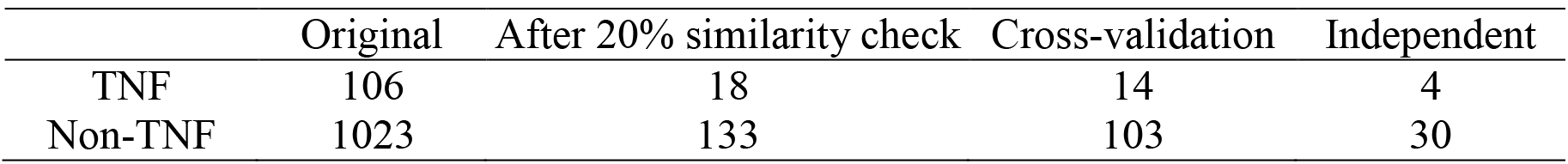
Statistics of the surveyed TNF and non-TNF sequences.

|  | Original | After 20% similarity check | Cross-validation | Independent |
| --- | --- | --- | --- | --- |
| TNF | 106 | 18 | 14 | 4 |
| Non-TNF | 1023 | 133 | 103 | 30 |

In order to make our analysis obvious and replicable, we provide our dataset at http://bio216.bioinfo.yzu.edu.tw/TNFPred.

### Feature extraction for identifying tumor necrosis factors

According to the fundamental concepts, in our research, every protein sequence is divided into segments with same length and allowed overlapping. We then trained the neural network via FastText [40-42] to generate the word embedding vector corresponding to each word. During this method, rather than employing a distinct vector illustration for every word, we considered inner arrangement in every word: each word w was described as a bag of character n-grams. In this manner, particular boundary characters “<“ and “>“ are supplementary at the start and finish of words. If taking QIGEF an instance, the following character n-grams: “<QI”, “QIG”, “IGE”, “GEF”, “EF>” and the particular sequence “<QIGEF >” will be counted when n = 3. For this reason, content of shorter words may be retained, which may seems as as *n*-grams of different words. This conjoint permits the meanings of suffixes/prefixes to be taken. This enable us to make the most of characteristics from rare words, that contributes considerably to the potency of the tactic [41]. It should be noted that we unbroken all the initial parameters of FastText and specify the embedding vector dimension as one. This implies that every biological word is delineated by only 1 real number.

Since surveyed sequence lengths are changing, the number of split words also varies. Machine learning algorithms, however, involve an equal amount of variables in the data samples. We addressed this difficulty using two steps: (1) First, we built L, an ordered list containing all protein words from the training data. (We v to denote the number of words in L), (2) Second, we used v embedding vectors appended one after another to describe each data sample. In this representation, the i^th^ vector is the embedding vector ensemble to the i^th^ word within the ordered list. During this step, it is important to note that if the i^th^ word is not present in the protein sequence, its corresponding embedding vector is adequate to zero. Likewise, if the i^th^ word emerges m times within the sequence, its corresponding embedding vector was increased m times. In this fashion, we would quantify the prevalence or contribution of the every biological word to a full feature vector. These biological words can be correlated to protein sequence motifs, that show their valuable characteristics for discriminating protein function [43]. The use of these characteristics enabled us to gain a lot of discriminatory features for more efficient prediction. The flow chart in Figure2 depicts our extraction technique for the case with a pair of sequences and biological word lengths equal to 3. In Additional file 1, we present a more detailed step-by-step demonstration of our feature extraction method with more sample sequences and selected n-gram length of 3. Moreover, rather than using fixed n-grams, with n are segmentation lengths of 1, 2, 3, 4 and 5, we conducted several mixtures among these grams, where we expected to find the best merge among different feature types to produce truly helpful features. The total number of these hybrid features is accumulated by surveyed grams.

**Figure 2.**
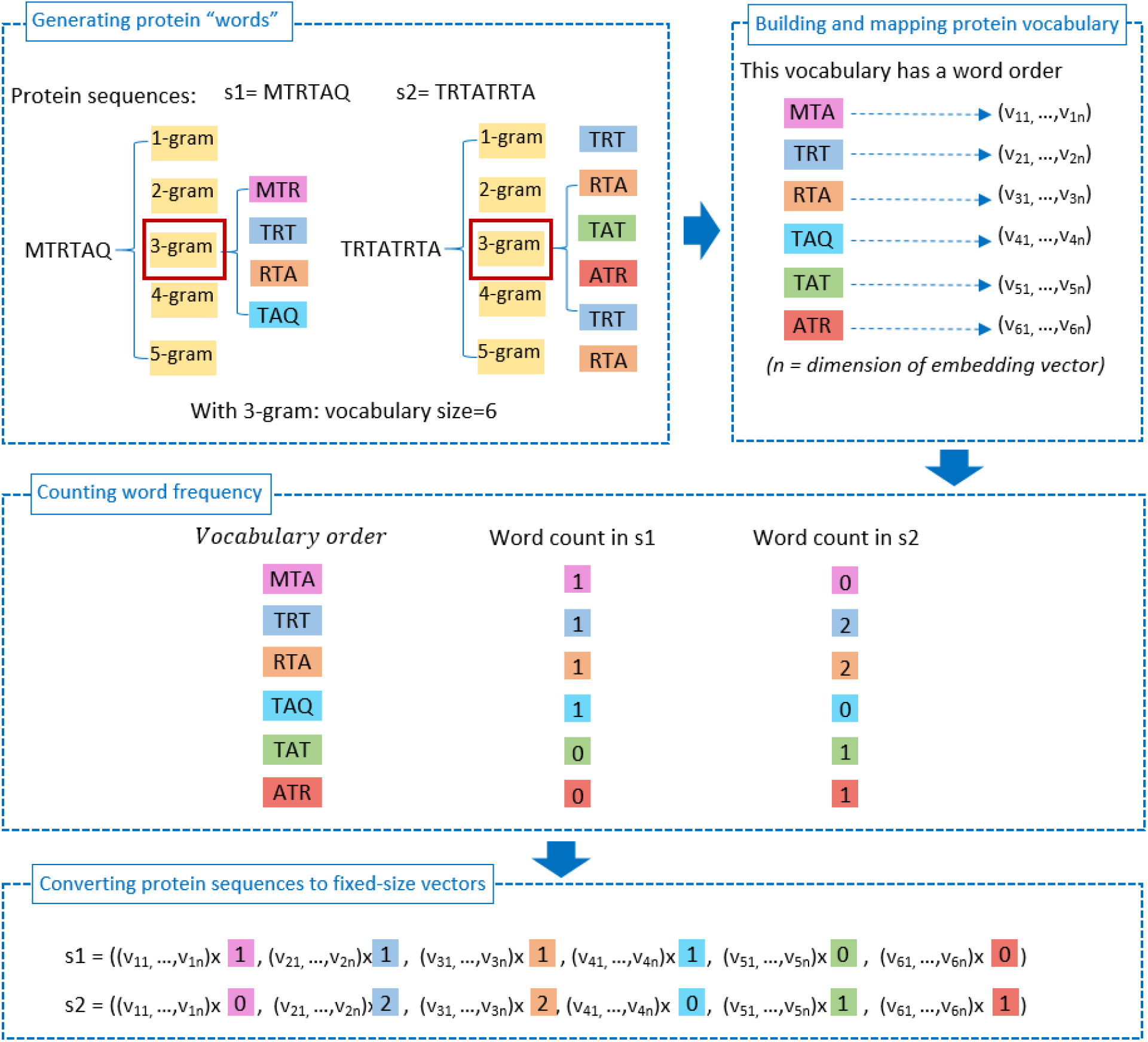
The 4-step flowchart illustrating our approach of utilizing word embedding vectors to create feature sets. In this illustration, there are 2 sequences and segmentation size is equal to 3.

### Assessment of predictive ability

Our study is a binary classification problem. First, we employed a five-fold cross-validation technique to develop and evaluate our models during the training process. Next, we assessed the capacity of our models in predicting unseen data using the independent dataset. We used four widely-used metrics sensitivity (Sen), specificity (Spec), accuracy (Acc) and Matthews’s correlation coefficient (MCC) (see, e.g. [28, 44-47]) to measure the predictive performance. In Additional file 2, the formulae of these metrics are presented. When a single metric is required to assess a predictive model globally, we used the area under the receiver operating characteristic (ROC) curves (AUC) score [48, 49].

## Results

### Number of features with n-gram sizes

It is necessary to figure out the quality as well as the quantity of extracted features for the learning process. Therefore, we calculated the number of features of word embedding-based vectors that were input in our binary classifiers (see Table 2). As we randomly divided the data into the training part and testing part and repeated this process for 10 times resulting 10 datasets for experiments, the numbers of n-grams vary from dataset to dataset. We found that when biological word length was equal to 3 (n=3), the numbers of features were highest (from 1736 to 1915) compared to those with length 1, 2, 4, or 5. In this group, the numbers of features when the length of biological words is equal to 2 is the second highest (from 395 to 398). Therefore, when we combined between 2-gram and 3-gram features, the total number of hybrid features is the highest one (from 2131 to 2313).

**Table 2.** Number of features input in our binary classifiers.

| Feature types | Number of features |
| --- | --- |
| 1-gram | 20 |
| 2-gram | 395 → 398 |
| 3-gram | 1736 → 1915 |
| 4-gram | 60 → 83 |
| 5-gram | 6 → 11 |
| 1-gram and 2-gram combined | 415 → 418 |
| 1-gram and 3-gram combined | 1756 → 1935 |
| 2-gram and 3-gram combined | 2131 → 2313 |

### Amino acid composition analysis of TNFs and non-TNFs

We carried out statistical tests on the entire dataset to evaluate the distinction between TNFs and non-TNFs amino acid composition variance. We performed the tests on 3 types of composition: single amino acid, dipeptide and tripeptide. First, F-test has been used to test the null hypothesis that two datasets’ variances are equal. We presumed that there are equal variances between TNFs and non-TNFs. After running the test, in all three cases, we acknowledged F < F-critical, (the F and F-critical values of 3 cases are provided in the Table S1, Aditional file 0), so we accepted the null hypothesis. It implies that the variances of two datasets in terms of single amino acid, dipeptide and tripeptide composition are equal or the amino acid structures of the dataset are not substantially different. Therefore, an efficient approach for extracting useful features is critically needed to identify TNFs with high performance.

Next, we performed the unpaired t-test with values of amino acid, dipeptide and tripeptide composition on positive group (18 TNF sequences) and negative group (non-TNF sequences). The table S2, S3, S4 in the Additional file 0 displays the p-values of the tests.

From the table S2, we can see that the p-values are generally high which means that there are not much difference in amino acid compostion between TNF and non-TNF sequences. However, at the common criterion (p-value of 0.05), six amino acids C, G, K, W, Y, and V shows significant frequency difference. In addition, from the table S3, it can be seen that GL, LY, FG are amino acid pairs with lowest p-values which means that they may contribute more for distinguishing positive and negative data. It is also noted from the Table S4 that these patterns can be found in tripeptides with lowest p-values (GLY, FFG, FGA, LYY). The other tripeptides with lowest p-values are QDG, TFF and VYV.

### Visualization of word embedding-based features in representation space

In this analysis, we would like to envisage the surveyed protein feature vectors, which have a high number of dimensions and are very hard to see in the raw form. We performed t-Distributed Stochastic Neighbor Embedding [50] (t-SNE), an unsupervised, non-linear method mainly used for data exploration and high-dimensional visualization, on the whole dataset. t-SNE allows us to reduce data with hundreds or even thousands of dimensions to just two (forming a point in twodimensional space) and thus being suitable to plot our data on a two-dimensional space. We used the word embedding-based vectors generated from extraction method described in section 2.2 as input into t-SNE algorithm to create two-dimensional “maps” for TNF and non-TNF sequences. We also noted that t-SNE contains a tunable parameter, “perplexity”, which indicated balance attention between local and global aspects of visualized data. The perplexity value has a complex effect on the resulting pictures and typical values are between 5 and 50. In order to get a multiple view about our data, we performed our visualization with perplexity values of 25 and 50. Figure S1, S2, S3, S4, S5, S6, S7, S8 in Additional file 3 visualize our dataset corresponding to 5 different word lengths (length = 1, 2, 3, 4, 5) and 3 ways we combined n-gram features, respectively. From these figures, it is interesting that the two-dimensional “maps” for TNF and non-TNF sequences have different shapes. Moreover, features with n-gram, where n takes values of 1, 2, 4, and 5 are points that tend to be farther from each other compared to those of the rest. In addition, when we changed the perplexity from 25 to 50, the shape comprised from the points changed differently among feature types.

### The influence of n-gram sizes on the performance of the models

In this prior research, we intended to compare the efficacy of distinct segmentation sizes and the mixture of n-gram used. The feature sets were produced as outlined in section 2.2. We used SVM as the binary classifier. (We used scikit-learn library (version 0.19.1) for building all classifiers in this study). In both 5-fold cross-validation and independent test, we assessed the general performance of each experiment using the AUC scores. The outcomes with highlighted best values in bold are shown in Table 3. We found that among 5 segmentation sizes, feature type corresponding to biological words of length=3 helped yielded the highest average performance. This may come from the larger number of features because when the biological words length is equal to 3 (n=3), the numbers of features are highest (from 1736 to 1915, see Table 2). Furthermore, model with the 2-gram and 3-gram combined feature set achieved the best average AUC of 0.984±0.298 and 0.998±0.277 on cross-validation and independent test, respectively. From these outcomes, we chose this feature type for further experiments.

**Table 3.** AUC performance of SVM classifier on embedding features with different biological word lengths.

| Feature | AUC |  |
| --- | --- | --- |
|  | 5-fold cross-validation | Independent |
| 1-gram | 0.856±0.485 | 0.848±0.525 |
| 2-gram | 0.901±0.416 | 0.883±0.599 |
| 3-gram | 0.934±0.423 | 1±0 |
| 4-gram | 0.617±0.63 | 0.563±0.751 |
| 5-gram | 0.543±0.539 | 0.574±0.686 |
| 1-gram and 2-gram combined | 0.952±0.416 | 0.934±0.48 |
| 1-gram and 3-gram combined | 0.96±0.42 | 0.921±0.497 |
| 2-gram and 3-gram combined | **0.984±0.298** | **0.998±0.277** |

*Note:* (Each result is reported in format: $m \pm d$ , where $m$ is the mean and $d$ is the standard deviation across the ten runs)

### Comparison between proposed features and other sequence-based feature types

We used SVM to evaluate the efficiency of our suggested features against 7 frequently used feature types i.e. amino acid composition (AAC), amino acid pair composition (DPC), position-specific scoring matrix (PSSM) features and the combination of these types. After a search for the ideal parameters, we conducted these experiments on both cross-validation and independent data. We found that with our suggested features, the SVM models reached the best performance (95.82%, 97.59%, 83.67%, 0.83 on the validation data, 96.49%, 98%, 85%, 0.86 on the independent data in terms of accuracy, specificity, sensitivity, and MCC, respectively, see Table 4). These values demonstrate that, compared to other sequence-based feature kinds, our word embedding-based features are more discriminatory and efficient. For long time, it is well-known that evolutionary information or PSSM is an efficient feature type to solve numerous bioinformatics problem yet it takes a lot of time to be created. These results show that, with our approach for generating features, we can save much time on feature extraction phase.

**Table 4.**
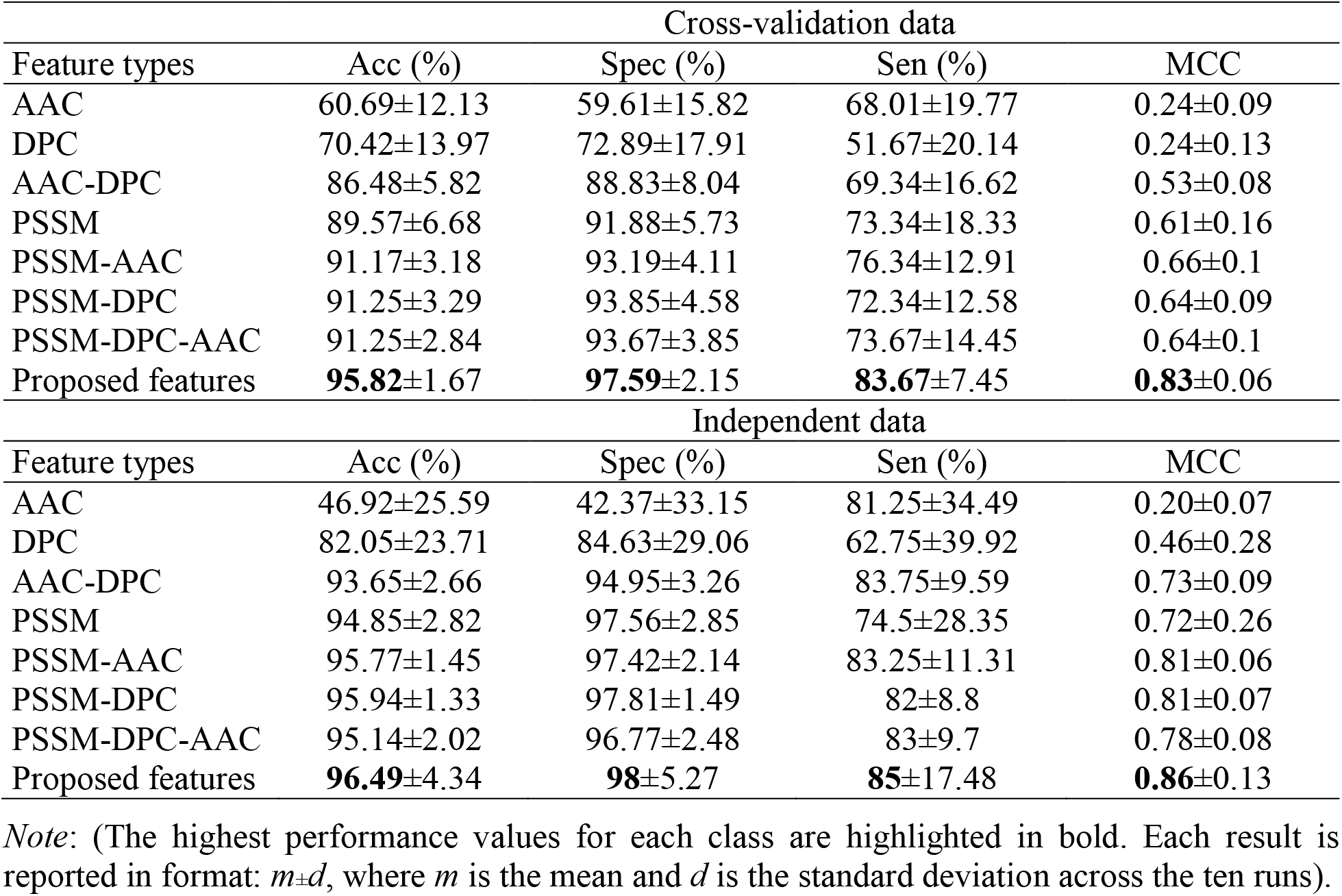
Performance comparison of proposed features with AAC, DPC, PSSM, and the combined features.

| Feature types | Cross-validation data |  |  |  |
| --- | --- | --- | --- | --- |
|  | Acc (%) | Spec (%) | Sen (%) | MCC |
| AAC | 60.69±12.13 | 59.61±15.82 | 68.01±19.77 | 0.24±0.09 |
| DPC | 70.42±13.97 | 72.89±17.91 | 51.67±20.14 | 0.24±0.13 |
| AAC-DPC | 86.48±5.82 | 88.83±8.04 | 69.34±16.62 | 0.53±0.08 |
| PSSM | 89.57±6.68 | 91.88±5.73 | 73.34±18.33 | 0.61±0.16 |
| PSSM-AAC | 91.17±3.18 | 93.19±4.11 | 76.34±12.91 | 0.66±0.1 |
| PSSM-DPC | 91.25±3.29 | 93.85±4.58 | 72.34±12.58 | 0.64±0.09 |
| PSSM-DPC-AAC | 91.25±2.84 | 93.67±3.85 | 73.67±14.45 | 0.64±0.1 |
| Proposed features | **95.82±1.67** | **97.59±2.15** | **83.67±7.45** | **0.83±0.06** |
| Feature types | Independent data |  |  |  |
|  | Acc (%) | Spec (%) | Sen (%) | MCC |
| AAC | 46.92±25.59 | 42.37±33.15 | 81.25±34.49 | 0.20±0.07 |
| DPC | 82.05±23.71 | 84.63±29.06 | 62.75±39.92 | 0.46±0.28 |
| AAC-DPC | 93.65±2.66 | 94.95±3.26 | 83.75±9.59 | 0.73±0.09 |
| PSSM | 94.85±2.82 | 97.56±2.85 | 74.5±28.35 | 0.72±0.26 |
| PSSM-AAC | 95.77±1.45 | 97.42±2.14 | 83.25±11.31 | 0.81±0.06 |
| PSSM-DPC | 95.94±1.33 | 97.81±1.49 | 82±8.8 | 0.81±0.07 |
| PSSM-DPC-AAC | 95.14±2.02 | 96.77±2.48 | 83±9.7 | 0.78±0.08 |
| Proposed features | **96.49±4.34** | **98±5.27** | **85±17.48** | **0.86±0.13** |

*Note:* (The highest performance values for each class are highlighted in bold. Each result is reported in format: $m \pm d$ , where $m$ is the mean and $d$ is the standard deviation across the ten runs).

### Comparison of different advanced classifiers using embedding features

We used the feature kind selected from the results of the experiments described in section 3.4 as inputs of 5 commonly used machine learning algorithms namely Support Vector Machine [51], k Nearest Neighbor (kNN) [52], RandomForest (RF) [53], Naïve Bayes [54] and QuickRBF [55-58]. We used cross-validation data and independent data for these experiments. We also searched for the best parameters for each algorithm. The aim of this assessment is to find out which classifier obtains the greatest outputs given this kind of feature. Table 5 shows the outcomes with the greatest average performance values highlighted in bold. We discovered that, the SVM classifier outperformed the other classifiers on both cross-validation and independent data on the same suggested dataset in terms of AUC (see Table 5). Accordingly, we employed SVM to build the final prediction model.

**Table 5.**
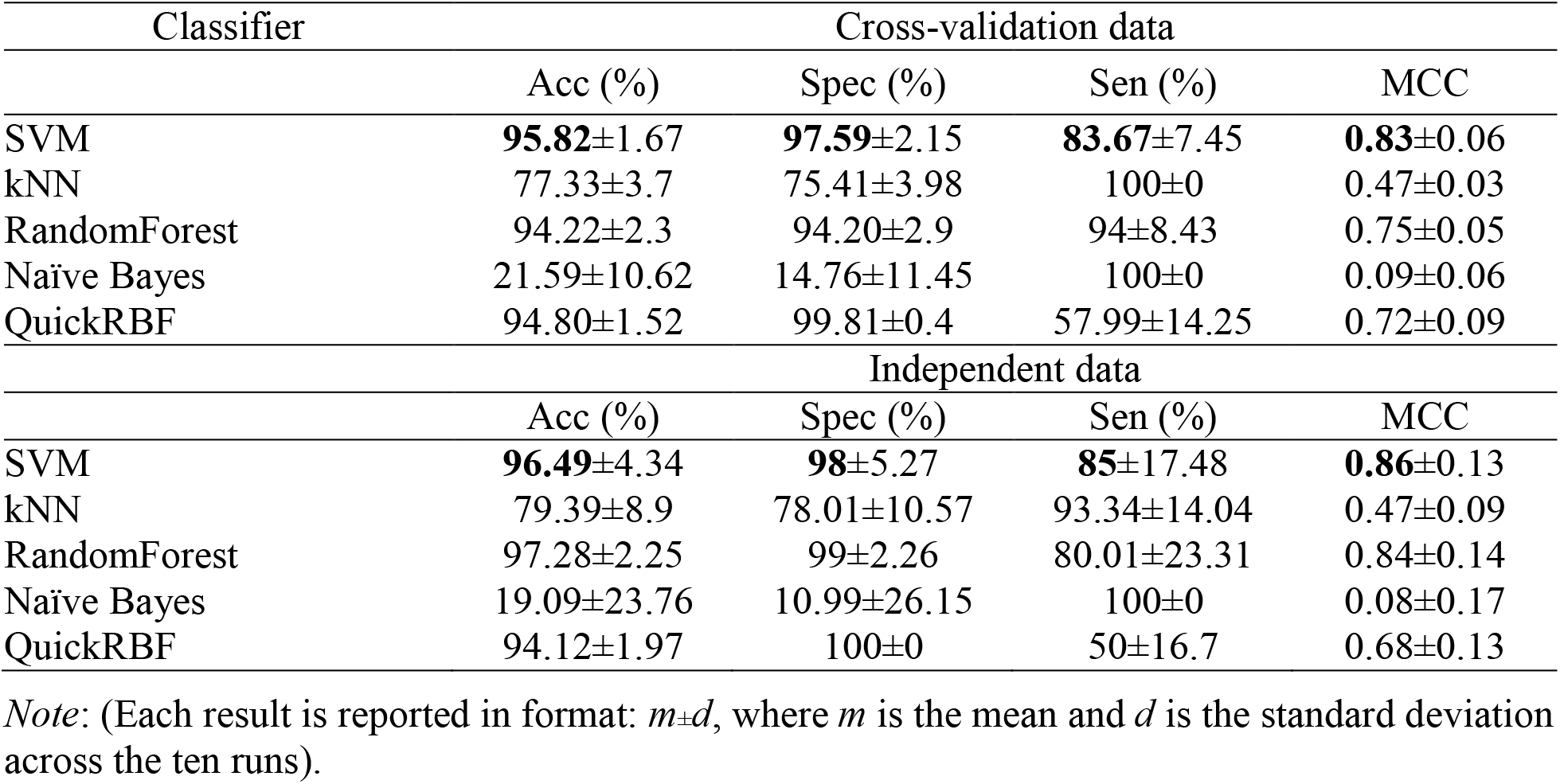
Performance comparison of five commonly used binary classifiers on proposed features

| Classifier | Cross-validation data |  |  |  |
| --- | --- | --- | --- | --- |
|  | Acc (%) | Spec (%) | Sen (%) | MCC |
| SVM | **95.82±1.67** | **97.59±2.15** | **83.67±7.45** | **0.83±0.06** |
| kNN | 77.33±3.7 | 75.41±3.98 | 100±0 | 0.47±0.03 |
| RandomForest | 94.22±2.3 | 94.20±2.9 | 94±8.43 | 0.75±0.05 |
| Naïve Bayes | 21.59±10.62 | 14.76±11.45 | 100±0 | 0.09±0.06 |
| QuickRBF | 94.80±1.52 | 99.81±0.4 | 57.99±14.25 | 0.72±0.09 |
| Classifier | Independent data |  |  |  |
|  | Acc (%) | Spec (%) | Sen (%) | MCC |
| SVM | **96.49±4.34** | **98±5.27** | **85±17.48** | **0.86±0.13** |
| kNN | 79.39±8.9 | 78.01±10.57 | 93.34±14.04 | 0.47±0.09 |
| RandomForest | 97.28±2.25 | 99±2.26 | 80.01±23.31 | 0.84±0.14 |
| Naïve Bayes | 19.09±23.76 | 10.99±26.15 | 100±0 | 0.08±0.17 |
| QuickRBF | 94.12±1.97 | 100±0 | 50±16.7 | 0.68±0.13 |

*Note:* (Each result is reported in format: $m \pm d$ , where $m$ is the mean and $d$ is the standard deviation across the ten runs).

### Classification on the web

We created a convenient and publicly accessible web server called TNFPred to illustrate our work. TNFPred is the web server especially trained for identifying tumor necrosis factors from other cytokines using support vector machine algorithm and proposed hybrid word embedding-based features. The user can use http://bio216.bioinfo.yzu.edu.tw/tnfpred to submit the sequences and assess our technique. Biologists are able to determine whether a cytokine is a tumor necrosis factor or not based on its amino acid sequence, according to this server.

## Discussions

In this work, we provided a computational model for classifying tumor necrosis factors from cytokines. We supported biologists with data for their experiment replications and scholarly work with reliable cytokine and non-cytokine sequence information. Additionally, our study was carefully-designed to ensure the reliability of the predictive model because we exactingly adhered to the 5-step rule proposed by Chou et al. We also carried the experiments with 10 runs, each with the same data but different training and testing data partitions to get more insight about the dataset. Although we kept the same sequence number distributions over these two parts, it is interesting to find that the performance is influenced by the random partitioning of data into training and testing data parts. This is reflected in the standard deviation across 10 runs. In this regard, sensitivity scores are the most unpredictable ones. We think a reason for this instability may come from the much insufficiency of positive data samples (see Table 1). Additionally, with our data, in the case of 3-gram, 4-gram and 5-gram, the number of biological words generated (see Table 2) are far from reaching the maximum possible number following this formulae: (20) ^length^, where 20 is the number of standard amino acids and length is the size for sequence segmentation. This means that we did not have abundant samples to train the FastText model so that it can reach its best potential. Fortunately, when we accumulated 2-gram and 3-gram features to create a new feature set, we obtained the best average AUC and lowest standard deviation (see Table 2). In order to improve the performance as well as the stability of predictive models in future research, we suggest 2 tentative methods. First, for extracting useful features, scientists may try with another word representation approach such as contextualized word embeddings of which the same word (motif) will have different embeddings depending on its contextual use [59-61]. Second, for tackling the insufficience of positive data samples (e.g., TNF sequences in this study), transfer learning approach can be a good attempt [62]. As part of our future work, we are currently evaluating the effect of an optimal features based on contextualized word embeddings for comparison with the scheme used in this study. Moreover, we are also considering transfer learning approach for protein classification tasks with source data from proteins having general characteristics of cytokines, e.g., cell-signalling, and target data from specific kind of cytokines such as TNFs.

## Conclusions

In this research, we have consistently applied word embedding techniques for identifying tumor necrosis factors from other cytokines. We assessed our performance on 5-fold cross-validation and independent testing dataset with support vector machine classifier and optimal features generated from word embeddings. Our technique showed a five-fold cross-validation accuracy of 95.82% and MCC of 0.83 for predicting TNFs among cytokines. The accuracy of independent datasets is 96.49% and MCC is 0.96, respectively. This strategy strictly follows the guidelines of 5-step rule, which makes our predictor reliable compared to other published works. Our suggested approach is also simpler, less laborious and much quicker to generate feature sets. In addition, our work could provide a foundation for future studies that can utilize natural language processing tactics in bioinformatics and computational biology.

## Declaration

## List of abbreviations

AAC: amino acid composition
Acc: accuracy
AUC: Area Under The Curve
DPC: dipeptide composition
IFN: interferon
IL: interleukin
kNN: k-nearest neighbor
MCC: Matthew’s correlation coefficient
NGF: neuron growth factor
NLP: natural language processing
PSSM: position specific scoring matrix
RF: random forest
ROC: receiver operating characteristic
Sen: sensitivity
Spec: specificity
SVM: support vector machine
TGF-b: transforminggrowth factor b
TNF: tumor necrosis factor
TPC: tripeptide composition
t-SNE: t-Distributed Stochastic Neighbor Embedding

## Ethics approval and consent to participate

Not applicable

## Consent for publication

Not applicable

## Availability of data and material

The datasets generated and/or analyzed during the current study are available in the web server at http://bio216.bioinfo.yzu.edu.tw/tnfpred

## Authors’ contributions

Analyzed the data: TTDN DVP. Designed and performed the experiments: TTDN. Wrote the paper: TTDN NQKL. Building the web server: QTH. Read and approved the final version: TTDN NQKL QTH DVP YYO.

## Additional file 0

**Table S1:** F and F-critical values from F-test to test the null hypothesis that two datasets’ variances are equal in terms of single amino acid, dipeptide and tripeptide composition

|  | Amino acid | Dipeptide | Tripeptide |
| --- | --- | --- | --- |
| F | 1.063 | 0.489 | 0.892 |
| F Critical one-tail | 2.173 | 0.848 | 0.964 |

**Table S2:** p-values from the unpaired student T-test on single amino acid composition on positive group (18 TNF sequences) and negative group (non-TNF sequences) with the assumption that the variance are equal

|  |  |  |  |  |  |  |  |  |  |  |
| --- | --- | --- | --- | --- | --- | --- | --- | --- | --- | --- |
| Amino acid | A | R | N | D | C | Q | E | G | H | I |
| p-value | 0.434 | 0.229 | 0.402 | 0.410 | 0.008 | 0.289 | 0.414 | 0.005 | 0.492 | 0.124 |
| Amino acid | L | K | M | F | P | S | T | W | Y | V |
| p-value | 0.318 | 0.038 | 0.093 | 0.379 | 0.439 | 0.300 | 0.211 | 0.029 | 0.005 | 0.034 |

**Table S3:** Top 20 dipeptides with the lowest p-values from the unpaired student T-test on dipeptide composition on positive group (18 TNF sequences) and negative group (non-TNF sequences) with the assumption that the variance are equal

|  |  |  |  |  |  |  |  |
| --- | --- | --- | --- | --- | --- | --- | --- |
| Dipeptide | GL | LY | FG | SW | LV | EG | VY |
| p-value | 1.98E-08 | 1.07E-07 | 1.75E-07 | 3.29E-06 | 2.76E-05 | 2.76E-05 | 3.94E-05 |
| Dipeptide | QV | PW | QD | YF | GA | IY | FQ |
| p-value | 4.34E-05 | 6.59E-05 | 7.66E-05 | 3.60E-04 | 3.90E-04 | 6.24E-04 | 7.27E-04 |
| Dipeptide | HL | WE | YW | YY | AG | QI |  |
| p-value | 1.01E-03 | 1.03E-03 | 1.56E-03 | 1.86E-03 | 2.11E-03 | 2.35E-03 |  |

**Table S4:**
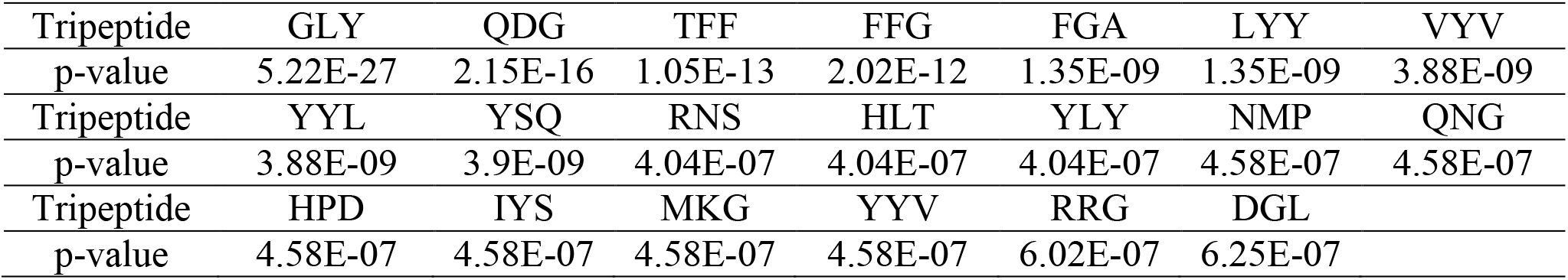
Top 20 tripeptides with the lowest p-values from the unpaired student T-test on tripeptide composition on positive group (18 TNF sequences) and negative group (non-TNF sequences) with the assumption that the variance are equal

|  |  |  |  |  |  |  |  |
| --- | --- | --- | --- | --- | --- | --- | --- |
| Tripeptide | GLY | QDG | TFF | FFG | FGA | LYY | VYV |
| p-value | 5.22E-27 | 2.15E-16 | 1.05E-13 | 2.02E-12 | 1.35E-09 | 1.35E-09 | 3.88E-09 |
| Tripeptide | YYL | YSQ | RNS | HLT | YLY | NMP | QNG |
| p-value | 3.88E-09 | 3.9E-09 | 4.04E-07 | 4.04E-07 | 4.04E-07 | 4.58E-07 | 4.58E-07 |
| Tripeptide | HPD | IYS | MKG | YYV | RRG | DGL |  |
| p-value | 4.58E-07 | 4.58E-07 | 4.58E-07 | 4.58E-07 | 6.02E-07 | 6.25E-07 |  |

## Additional file 1

Feature extraction method

A step-by-step demonstration of our feature extraction method with 3 sample sequences and selected n-gram length of 3

Split sequence into overlapping n-grams
| Sequence | Sequence content | Overlapping n-grams |
| --- | --- | --- |
| Seq1 | MTAASQANPYSI | → MTA **TAA AAS** ASQ SQA QAN ANP NPY PYS YSI (10 trigrams) |
| Seq2 | DTAASVTMSGAKGF | → DTA **TAA AAS** ASV SVT VTM TMS MSG SGA GAK AKG KGF (12 trigrams) |
| Seq3 | GNGKVHTQTAASNRFVAAS | → GNG NGK GKV KVH VHT HTQ **TAA AAS ASN** NRF RFV VAA **AAS** (13 trigrams) |

Using FastText to create (n-gram: one-dimension embedding vector) pairs
| No. | (Ngram→Embedding vector) pair | No. | (Ngram→Embedding vector) pair | No. | (Ngram→Embedding vector) pair |
| --- | --- | --- | --- | --- | --- |
| 1 | MTA→ (0.9891) | 11 | DTA→ (0.9546) | 21 | GNG→ (1.004) |
| 2 | TAA→ (1.0671) | 12 | ASV→ (0.8894) | 22 | NGK→ (0.9214) |
| 3 | AAS→ (1.0654) | 13 | SVT→ (0.8828) | 23 | GKV→ (0.9581) |
| 4 | ASQ→ (1.0096) | 14 | VTM→ (0.8964) | 24 | KVH→ (0.8148) |
| 5 | SQA→ (1.0147) | 15 | TMS→ (1.0257) | 25 | VHT→ (0.9581) |
| 6 | QAN→ (0.9790) | 16 | MSG→ (0.8317) | 26 | HTQ→ (0.9097) |
| 7 | ANP→ (1.0337) | 17 | SGA→ (0.9261) | 27 | ASN→ (0.9905) |
| 8 | NPY→ (0.8685) | 18 | GAK→ (0.9033) | 28 | NRF→ ( 0.9155) |
| 9 | PYS→ (0.9546) | 19 | AKG→ (0.8597) | 29 | RFV→ ( 0.9113) |
| 10 | YSI→ (0.8774) | 20 | KGF→ (0.9331) | 30 | VAA→ (0.8989) |

| Position in vocabulary | 1 | 2 | 3 | 4 | 5 | 6 | 7 | 8 | 9 | 10 | 11 | 12 | 13 | 14 | 15 | 16 | 17 | 18 | 19 | 20 | 21 | 22 | 23 | 24 | 25 | 26 | 27 | 28 | 29 | 30 |
| --- | --- | --- | --- | --- | --- | --- | --- | --- | --- | --- | --- | --- | --- | --- | --- | --- | --- | --- | --- | --- | --- | --- | --- | --- | --- | --- | --- | --- | --- | --- |
| 3-gram | MTA | TAA | AAS | ASQ | SQA | QAN | ANP | NPY | PYS | YSI | DTA | ASV | SVT | VTM | TMS | MSG | SGA | GAK | AKG | KGF | GNG | NGK | GKV | KVH | VHT | HTQ | ASN | NRF | RFV | VAA |
| Frequency of 3-grams in Seq1 | 1 | 1 | 1 | 1 | 1 | 1 | 1 | 1 | 1 | 1 | 0 | 0 | 0 | 0 | 0 | 0 | 0 | 0 | 0 | 0 | 0 | 0 | 0 | 0 | 0 | 0 | 0 | 0 | 0 | 0 |
| Frequency of 3-grams in Seq2 | 0 | 0 | 0 | 0 | 0 | 0 | 0 | 0 | 0 | 0 | 1 | 1 | 1 | 1 | 1 | 1 | 1 | 1 | 1 | 1 | 1 | 1 | 0 | 0 | 0 | 0 | 0 | 0 | 0 | 0 |
| Frequency of 3-grams in Seq3 | 0 | 1 | 2 | 0 | 0 | 0 | 0 | 0 | 0 | 0 | 0 | 0 | 0 | 0 | 0 | 0 | 0 | 0 | 0 | 0 | 1 | 1 | 1 | 1 | 1 | 1 | 1 | 1 | 1 | 1 |

The feature vector of each sequence is calculated as:

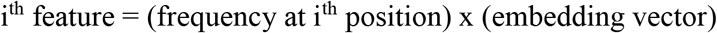

In our study, the dimension of the embedding vector is equal to 1, therefore the embedding vector is a scalar value.

Finally, the word embedding-based feature vectors are as below:

**<u>seq1:</u>** (0.9891, 1.0671, 1.0654, 1.0096, 1.0147, 0.9791, 1.0337, 0.8685, 0.9546, 0.8775, 0, 0, 0, 0, 0, 0, 0, 0, 0, 0, 0, 0, 0, 0, 0, 0, 0, 0, 0, 0)
**<u>seq2:</u>** (0, 0, 0, 0, 0, 0, 0, 0, 0, 0, 0.9546, 1.0671, 1.0654, 0.8895, 0.8829, 0.8965, 1.0257, 0.8318, 0.9262, 0.9034, 0.8598, 0.9332, 0, 0, 0, 0, 0, 0, 0, 0)
**<u>seq3:</u>** (0, 1.0671, 2.1308, 0, 0, 0, 0, 0, 0, 0, 0, 0, 0, 0, 0, 0, 0, 0, 0, 0, 1.0040, 0.9214, 0.9582, 0.8149, 0.9581, 0.9098, 0.9905, 0.9155, 0.9114, 0.8989)

These vectors were the input of binary classifiers.

## Additional file 2

Assessment of predictive ability

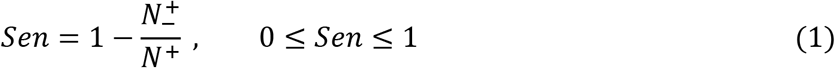

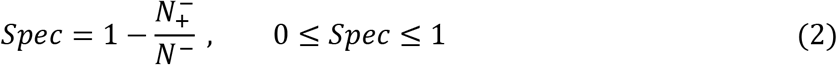

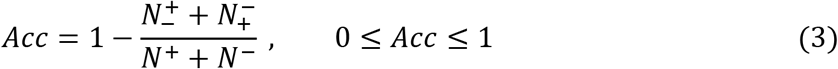

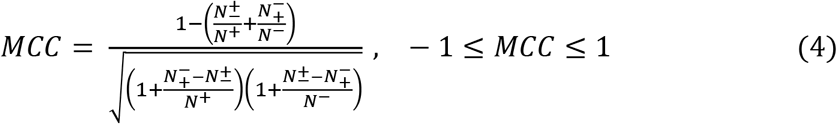

TP, FP, TN, FN stand for true positive, false positive, true negative, and false negative respectively.

The relations between these symbols and the symbols in Eqs. (1, 2, 3 and 4) are given by:

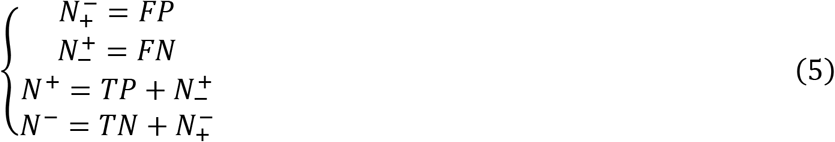

where, 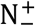 represents the number of positive samples (TNFs) incorrectly predicted to be negative samples (non-TNFs), 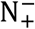 represents the number of negative samples incorrectly predicted to be positive sample, N^+^ represents the number of positive sample surveyed and N^−^ represents the number of negative sample surveyed.

## Additional file 3

Visualization of feature sets

**Figure S1a:**
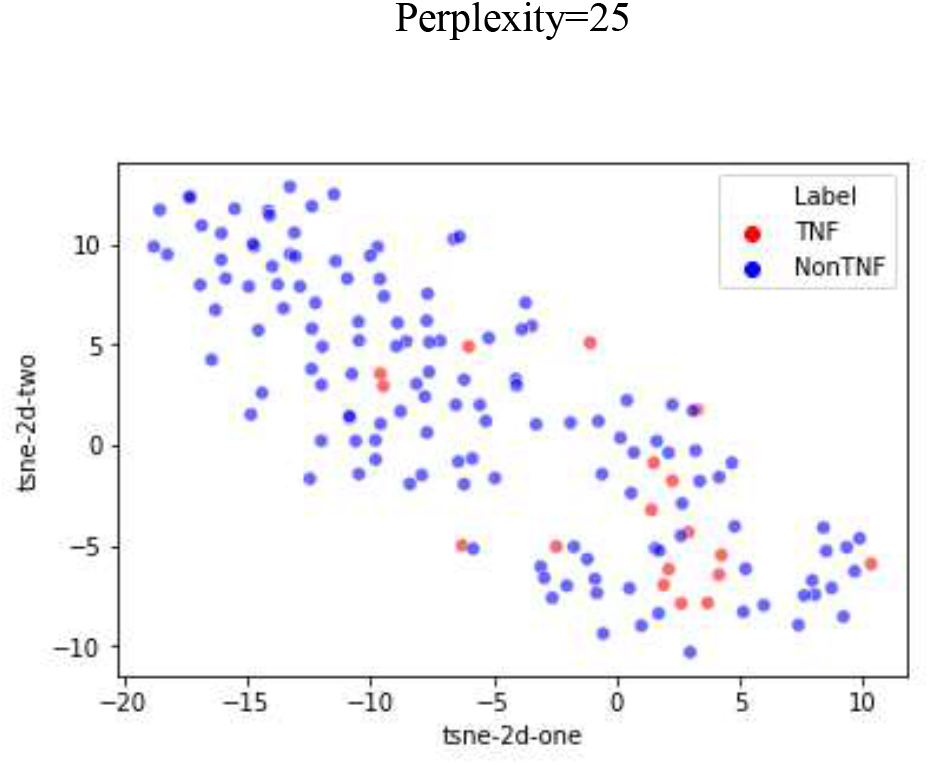
Visualization corresponding to protein feature vectors comprised from 1-gram embedding vectors with perplexity equal to 25

**Figure S1b:**
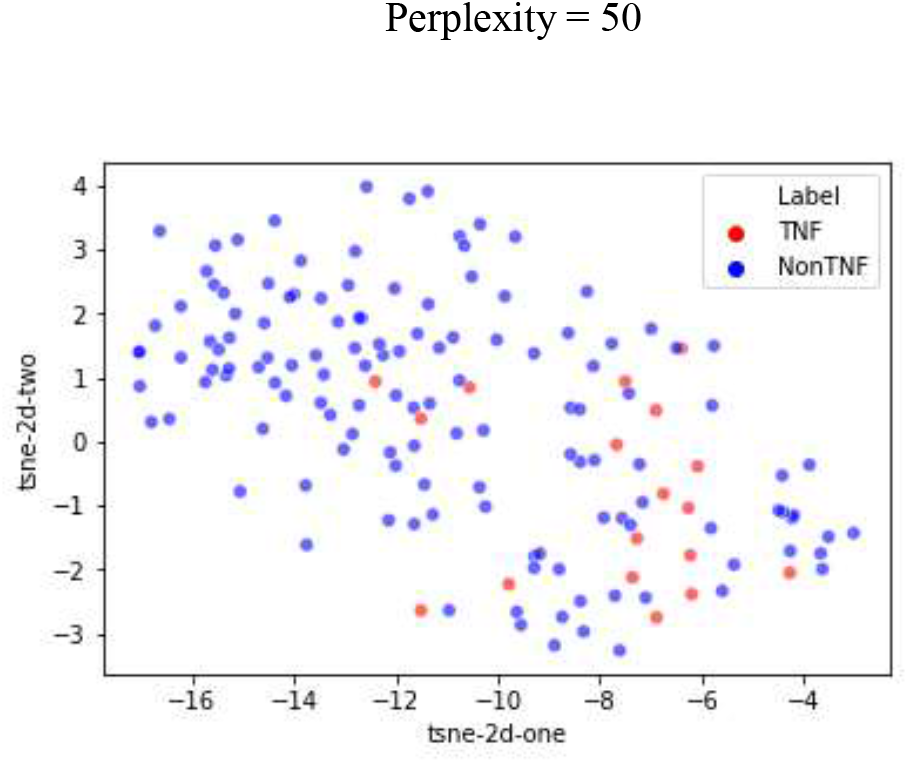
Visualization corresponding to protein feature vectors comprised from 1-gram embedding vectors with perplexity equal to 50

**Figure S2a:**
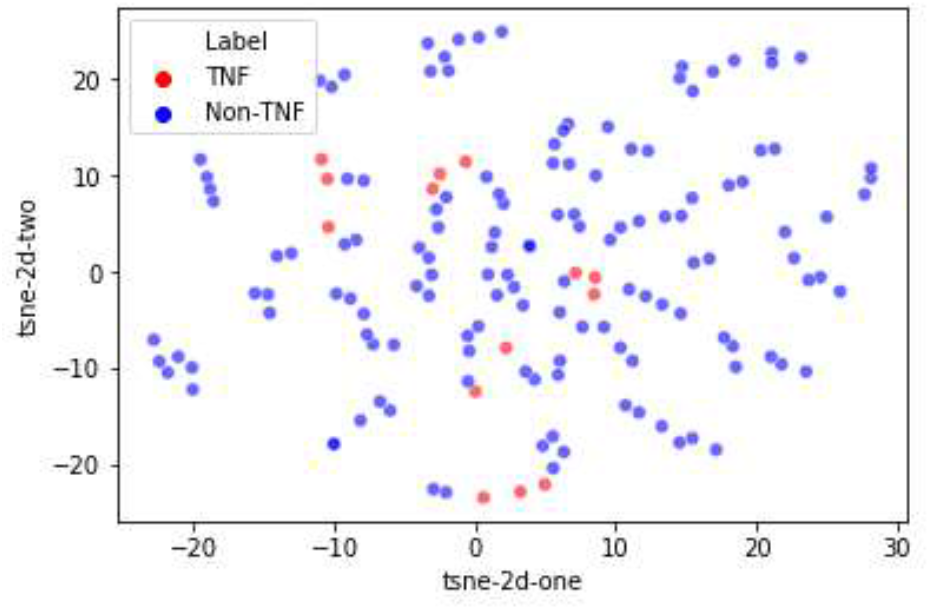
Visualization corresponding to protein feature vectors comprised from 2-gram embedding vectors with perplexity equal to 25

**Figure S2b:**
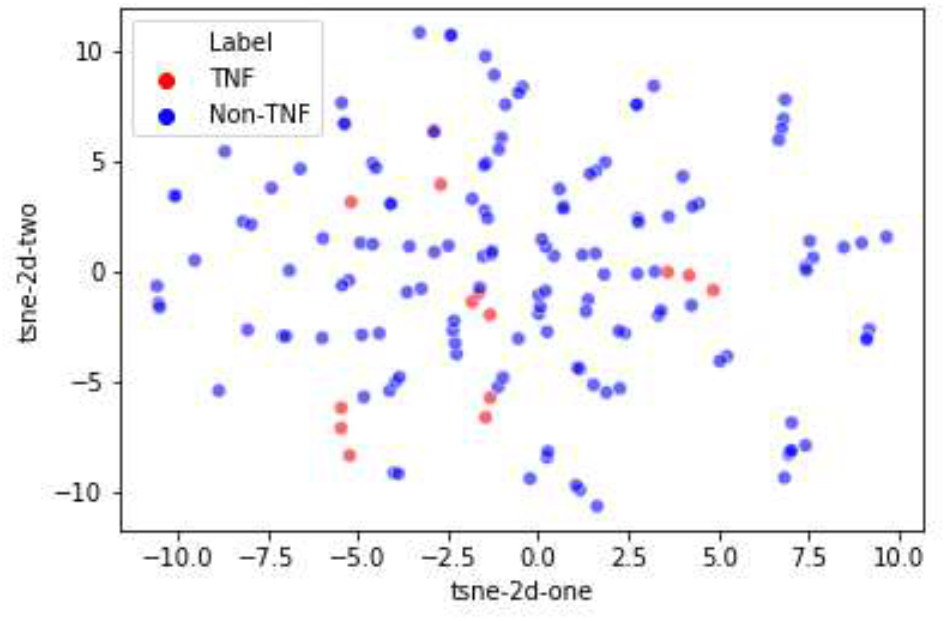
Visualization corresponding to protein feature vectors comprised from 2-gram embedding vectors with perplexity equal to 50

**Figure S3a:**
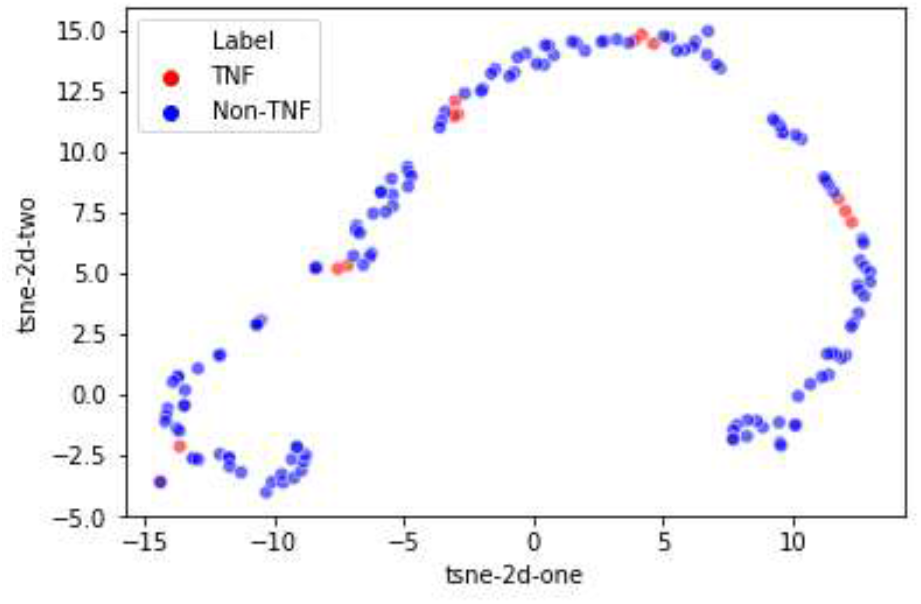
Visualization corresponding to protein feature vectors comprised from 3-gram embedding vectors with perplexity equal to 25

**Figure S3b:**
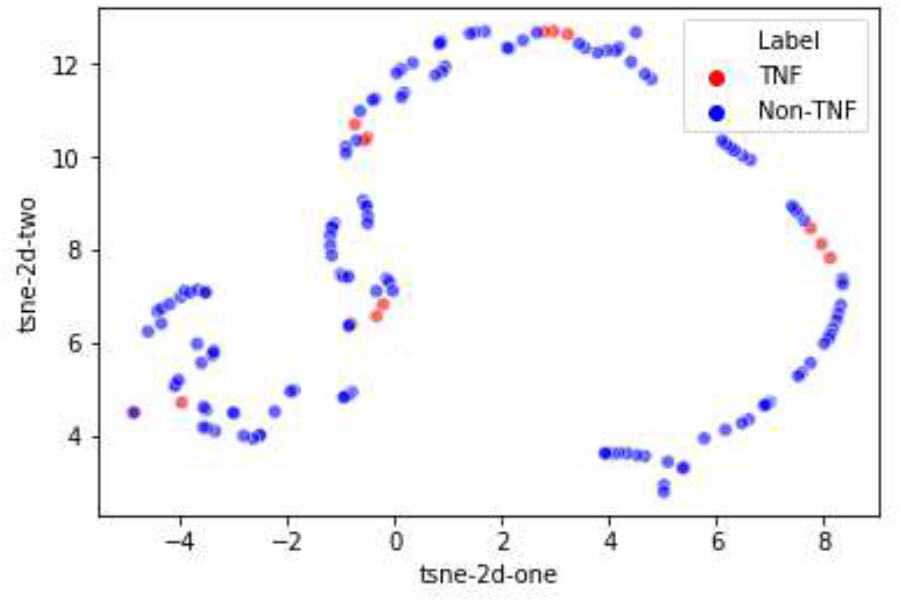
Visualization corresponding to protein feature vectors comprised from 3-gram embedding vectors with perplexity equal to 50

**Figure S4a:**
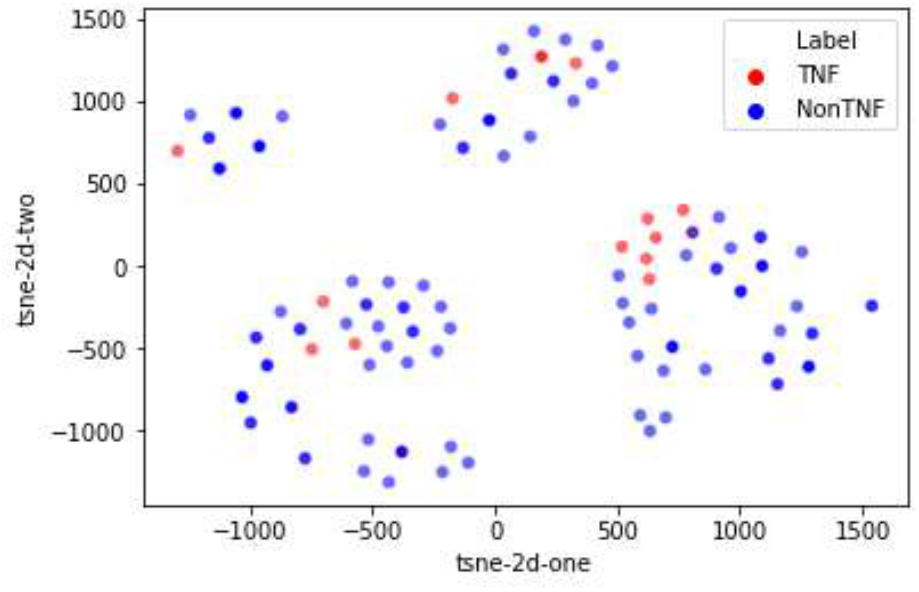
Visualization corresponding to protein feature vectors comprised from 4-gram embedding vectors with perplexity equal to 25

**Figure S4b:**
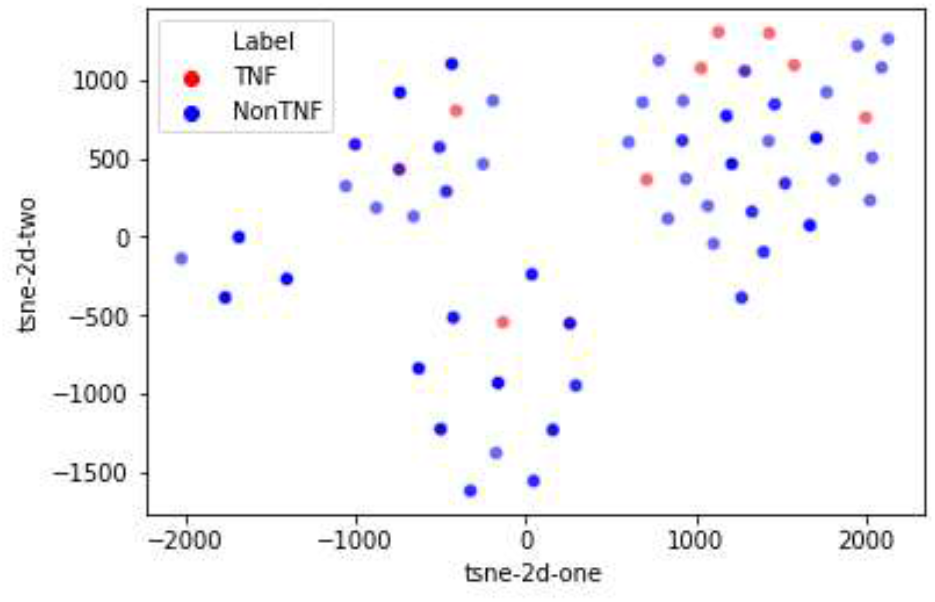
Visualization corresponding to protein feature vectors comprised from 4-gram embedding vectors with perplexity equal to 50

**Figure S5a:**
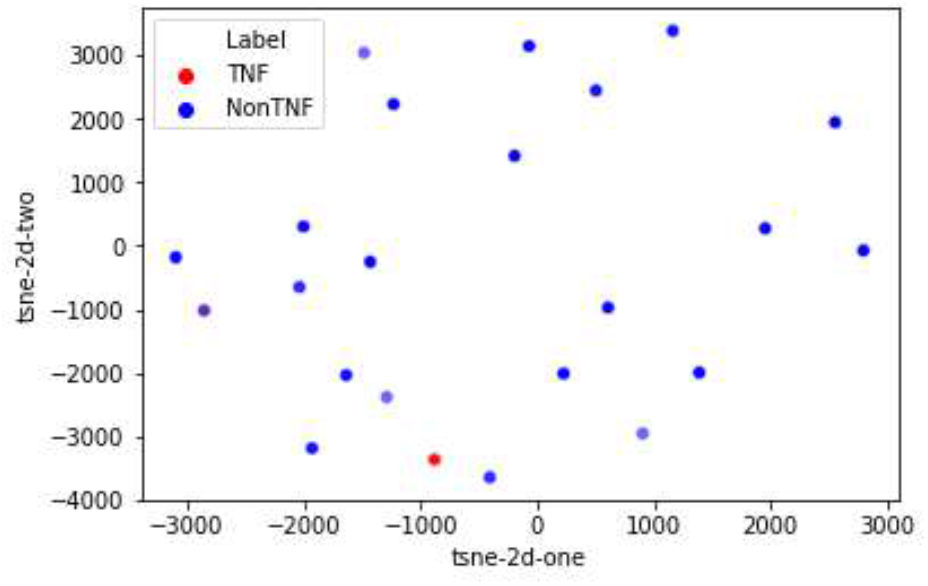
Visualization corresponding to protein feature vectors comprised from 5-gram embedding vectors with perplexity equal to 25

**Figure S5b:**
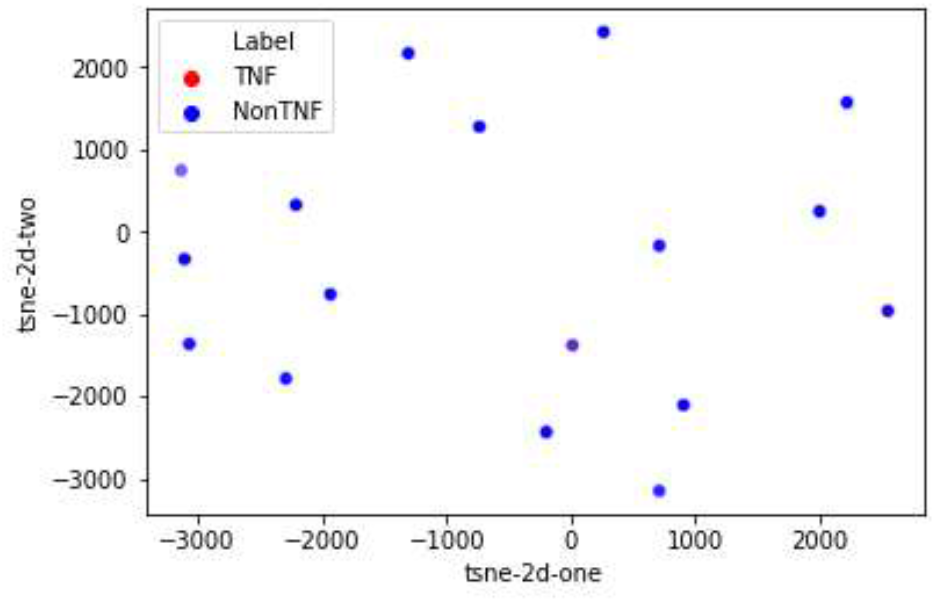
Visualization corresponding to protein feature vectors comprised from 5-gram embedding vectors with perplexity equal to 50

**Figure S6a:**
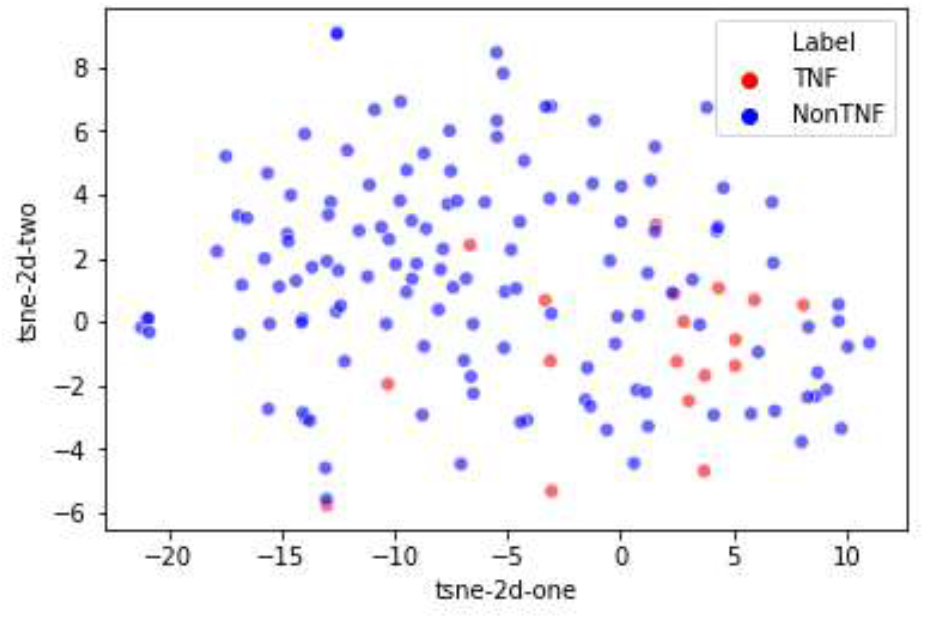
Visualization corresponding to protein feature vectors comprised from 1-gram and 2-gram embedding combined vectors with perplexity equal to 25

**Figure S6b:**
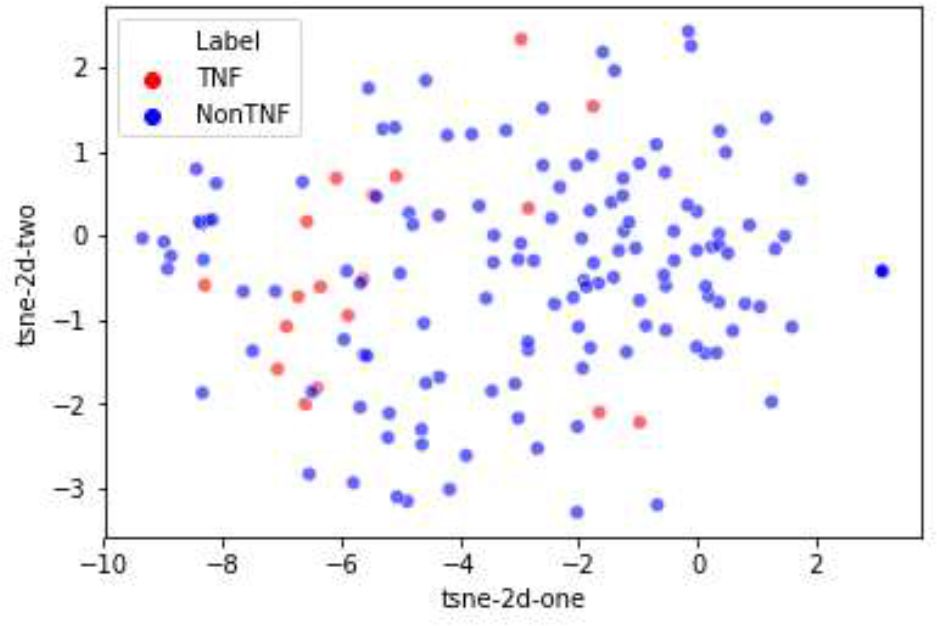
Visualization corresponding to protein feature vectors comprised from 1-gram and 2-gram embedding combined vectors with perplexity equal to 50

**Figure S7a:**
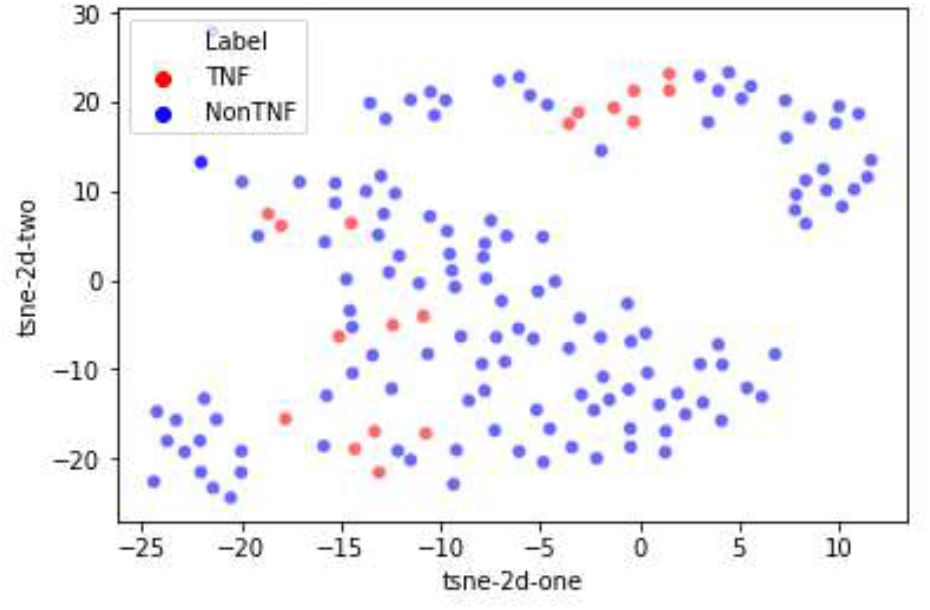
Visualization corresponding to protein feature vectors comprised from 1-gram and 3-gram embedding combined vectors with perplexity equal to 25

**Figure S7b:**
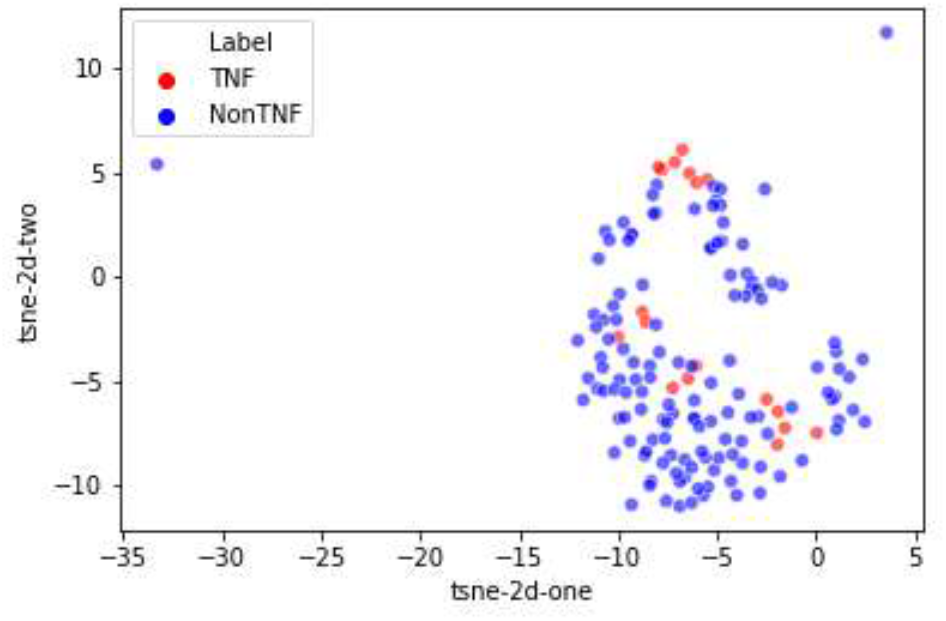
Visualization corresponding to protein feature vectors comprised from the combination of 2-gram and 3-gram embedding combined vectors with perplexity equal to 50

**Figure S8a:**
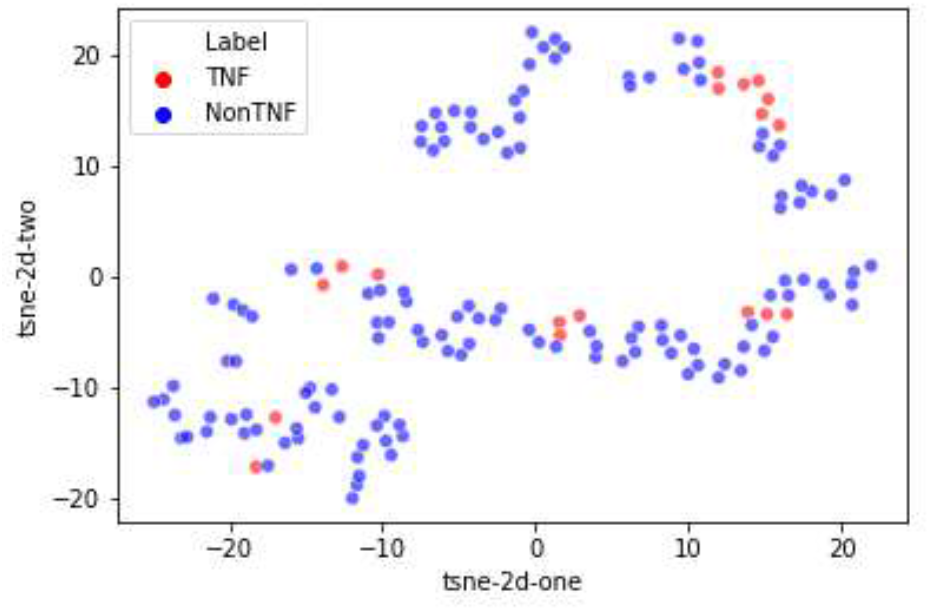
Visualization corresponding to protein feature vectors comprised from 2-gram and 3-gram embedding combined vectors with perplexity equal to 25

**Figure S8b:**
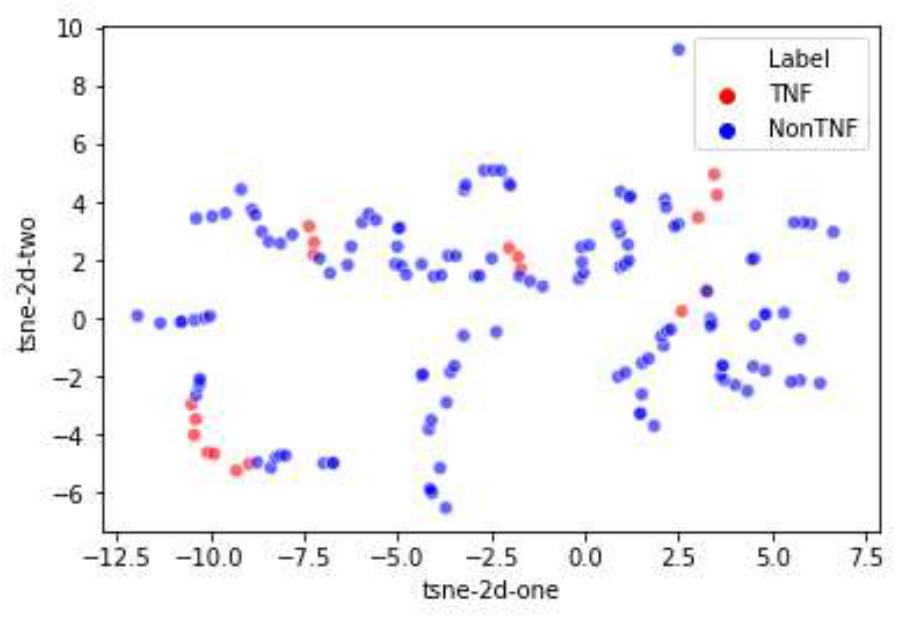
Visualization corresponding to protein feature vectors comprised from the combination of 2-gram and 3-gram embedding combined vectors with perplexity equal to 50

